# Deep learning and direct sequencing of labeled RNA captures transcriptome dynamics

**DOI:** 10.1101/2023.11.17.567581

**Authors:** Vlastimil Martinek, Jessica Martin, Cedric Belair, Matthew J Payea, Sulochan Malla, Panagiotis Alexiou, Manolis Maragkakis

## Abstract

Quantification of the dynamics of RNA metabolism is essential for understanding gene regulation in health and disease. Existing methods rely on metabolic labeling of nascent RNAs and physical separation or inference of labeling through PCR-generated mutations, followed by short-read sequencing. However, these methods are limited in their ability to identify transient decay intermediates or co-analyze RNA decay with cis-regulatory elements of RNA stability such as poly(A) tail length and modification status, at single molecule resolution. Here we use 5-ethynyl uridine (5EU) to label nascent RNA followed by direct RNA sequencing with nanopores. We developed RNAkinet, a deep convolutional and recurrent neural network that processes the electrical signal produced by nanopore sequencing to identify 5EU-labeled nascent RNA molecules. RNAkinet demonstrates generalizability to distinct cell types and organisms and reproducibly quantifies RNA kinetic parameters allowing the combined interrogation of RNA metabolism and cis-acting RNA regulatory elements.

## Introduction

The life cycle of RNA from transcription to decay is tightly regulated in cells. Unraveling of the kinetic parameters of RNA metabolism is critical for understanding gene regulation and cellular response to environmental cues. Existing methods for studying RNA dynamics in mammalian cells are based on short-read RNA sequencing (RNA-Seq) and involve the exposure of cells (pulse) to nucleoside analogs such as 4-thiouridine (4^SU^) or 5′-bromo-uridine (BrU) that get incorporated during transcription into newly synthesized RNAs ^1–8^. Subsequent steps involve either the chemical isolation of metabolically labeled RNAs before sequencing or the bioinformatic inference of labeling through PCR-generated mutations after sequencing ^9,10^. These methods have substantially simplified the methodology, but all share common limitations. Alternative splicing is a fundamental cellular process with biological and clinical relevance in which exons from the same gene are joined in different combinations, leading to different, but related, mRNA isoforms ^11^. However due to the limitation of using short-read sequencing, established methods cannot make high-confidence assignment of isoforms or collect information on individual RNA molecules. This prevents association of RNA kinetics with cis-acting transcriptional regulators such as alternative transcription start site and poly(A) site usage, post-transcriptional RNA modifications, and poly(A) tails, known regulators of RNA metabolism ^12^.

Nanopore sequencing has emerged as a promising avenue for the direct detection of nucleotides from the electrical current intensity as RNA molecules pass through nanopores ^13,14^. Besides canonical nucleotides, recent tools have been developed to also detect naturally occurring modified nucleotides. Some of these tools are comparative and require control samples to detect shifts in signal-based features that correlate with the presence of modifications ^15–18^ while others use training data to detect specific modifications in single samples ^19–23^. A recent work introduced nano-ID, a method that showed the feasibility to combine nanopore sequencing and machine learning to identify metabolically labeled RNAs in a single dRNA-Seq experiment following metabolic labeling with the nucleoside analog 5EU ^24^. However, our results show that nano-ID could not generalize to other experiments to be widely applicable.

In this work we present RNAkinet, a computationally efficient, convolutional, and recurrent neural network (NN) that identifies individual 5EU-modified RNA molecules following direct RNA-Seq (dRNA-Seq). RNAkinet can analyze entire experiments in hours, instead of days that nano-ID does, and predicts the modification status of RNA molecules directly from the raw nanopore signal without using basecalling or reference sequence alignment. We show that RNAkinet generalizes to sequences from unique experimental settings, cell types, and species and accurately quantifies RNA kinetic parameters, from single time point experiments. Being able to interrogate whole RNA molecules, RNAkinet allows, for the first time, the combined interrogation of RNA metabolism of single isoforms and cis-acting RNA regulatory elements, such as alternative splicing, poly(A) tail length and post-transcriptional RNA modifications.

## Results

### Data preparation and neural network design

To test nano-ID, we prepared data following labeling of HeLa cells with 5EU for 24 hours (h) as previously described ^24^. Given that the half-life of the majority of RNAs ranges in the order of a few hours, it is expected that the majority of RNA molecules would incorporate a 5EU ^1,25,26^. We defined this as our positive dataset while RNA from HeLa cells cultured without addition of 5EU served as our negative dataset (**Supplementary Table 1**). RNA from both datasets was sequenced directly on a MinION device as previously described ^24^. On our data, nano-ID required more than 7 days runtime and upwards of 800 Gb of memory use per sample on a high-performance system. Our results showed that nano-ID did not accurately distinguish labeled from unlabeled sequences, indicating that the nano-ID model had overfitted to the original training data (**Sup. Fig. 1A**).

Fully connected neural networks (NN) are subject to overfitting particularly when over-parameterized ^27^. Nano-ID comprises a fully connected neural network with 2,590,493 trainable parameters that include information from the raw electrical signal, basecalling and alignment that was trained on fewer than 700,000 examples (**Supplementary Table 2, bottom**). To address this gap, we aimed to develop a prediction tool, RNAkinet, that would a) accurately distinguish 5EU-labeled RNA, b) generalize to other datasets, c) significantly reduce the time and space requirements for execution, d) depend solely on the sequencer raw electrical signal to avoid dependencies on basecalling or alignment software, and e) allow robust quantification of the kinetics of RNA. RNAkinet consists of a NN that utilizes both convolutional ^28^ and recurrent layers to address the shortcomings of fully connected NN, aiming to reduce the parameter space ^29^ and integrate long- and short-range interactions between electrical signal in a read (**Fig. 1A**). We explored two distinct architecture designs (named RNAkinet and RNAkinet-TL, respectively) with the first encompassing a small convolutional neural network (**Fig. 1B**), while the second exploring transfer learning utilizing RODAN, a recently published RNA basecaller ^30^, as backbone (**Fig. 1C**). Transfer learning is used in order to fine-tune and repurpose the basecaller to detect 5EU ^31^. In both cases, a recurrent layer was placed atop the convolutional backbone. Finally, a dense layer was used to integrate the pooled result across an entire read to predict the probability of the read being labeled (**Fig. 1B, C**).

**Figure 1:**
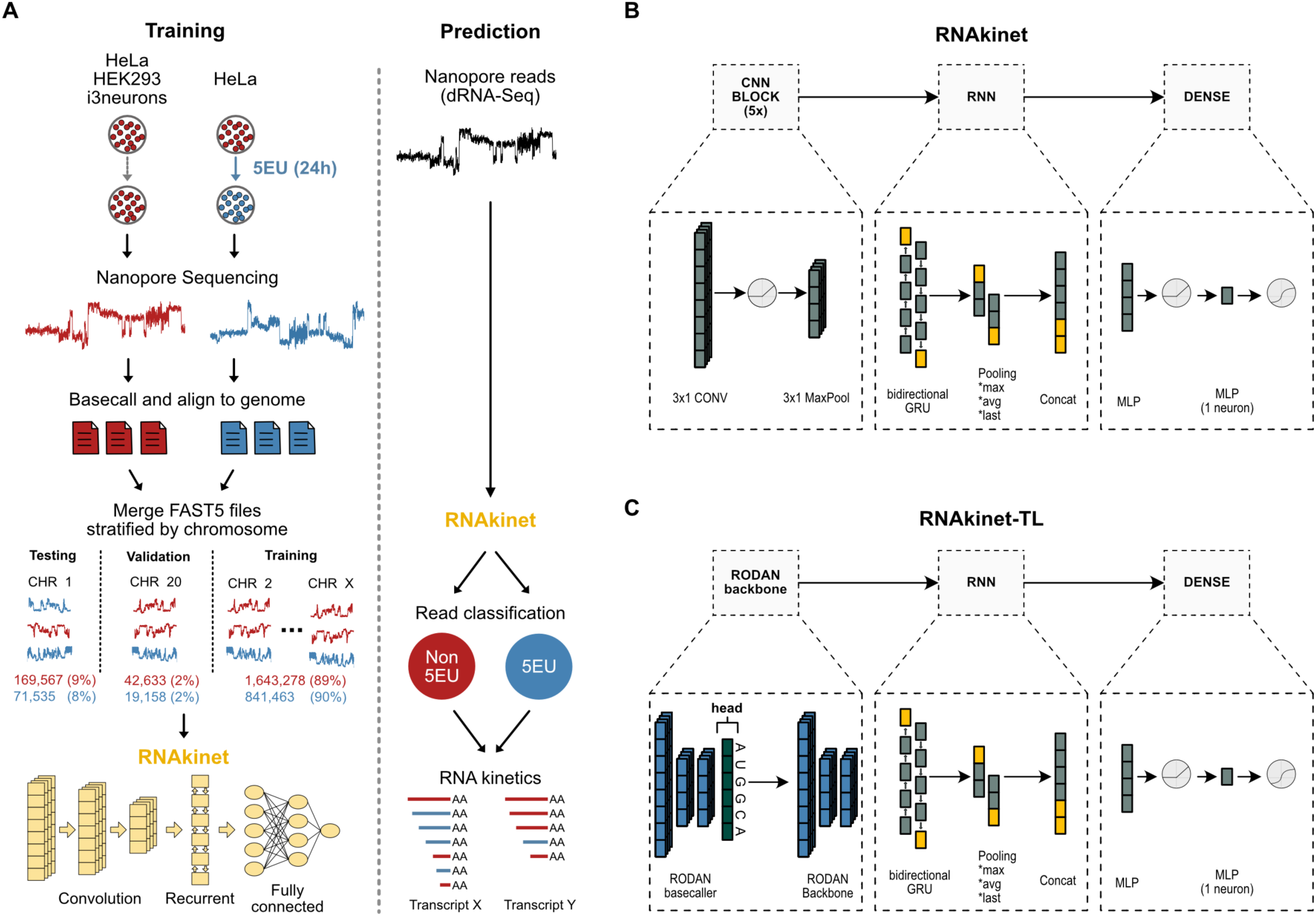
Data preparation and neural network design. **A)** Schematic of training and prediction workflow. Reads from chromosomes 1 and 20 were retained for testing and validation only. During prediction RNAkinet uses only the raw signal. **B-C**) Schematic representation of the architecture of RNAkinet (B) or an alternative architecture (RNAkinet-TL) involving transfer learning using the RODAN basecaller (C).

To ensure proper training, testing, and validation of RNAkinet, we followed a rigorous approach for data preparation. First, we performed basecalling and alignment to the human genome and subsequently separated the raw reads by chromosome. We isolated reads on chromosome 1 and 20, accounting for 8-9% and 2% of the total reads, to be used for testing and validation, respectively (**Fig. 1A**). This ensured that the network never interacted with sequences that exist in these two chromosomes during training, thus removing sequence level confounds from performance evaluation. Moreover, to enhance domain representation and to aid the NN into building a better representation of the experimental background and inherent nuisances in the data during training, we enriched our list of negative samples with data from independent dRNA-Seq experiments performed separately by different operators on distinct cell lines including HeLa, HEK293 cells and induced pluripotent stem cell-derived neurons (**Supplementary Table 1 and 2**). Overall, our training data aimed to reflect diversity among distinct experimental settings to enhance domain representation and model generalizability.

Raw electrical signal from the sequencing device is encoded in a one-dimensional array with varying size dependent on read length. Varying length input can be challenging for NNs and is typically addressed by padding to the length of the longest input. Given that in a typical dRNA-Seq experiment read lengths vary dramatically from a few bases to several kilobases in length (**Sup. Fig. 1B**), most sequences would need to be heavily padded. Processing of such signals would be computationally expensive, but more importantly, it would change the inherent semantics of continuous time series values of the electrical signal. For example, unlike pad tokens in Natural Language Processing networks ^32^, padding with zeros or any value would interfere with actual values in the electrical current that have a physical meaning in the signal generated from the sequencer. To avoid this, other tools have used fixed length windows of the input signal ^30,33^. However, this is not ideal for the purpose of detecting 5EU modification, since the 5EU labeling efficiency per nucleotide is low (2-3%) ^24^ raising the possibility of a labeled sequence not containing the modification in a given window. To address this, we trained the network on single whole sequences without padding and designed our network to accept sequences of any length and share learned features across the entire sequence.

Our training process involved randomly sampling from our positive and negative sets uniformly to ensure a balanced ratio for training. Since we used multiple datasets as negatives, those were sampled proportionally based on the number of reads they contained. Due to the large size of our dataset, we adopted a multi-read fast5 file format for quick loading, and we only loaded files when required to reduce the memory footprint. The same approach was used for inference, which ensured the network used the minimum amount of computational resources. Finally, we limited the input of the model to raw nanopore signal, not utilizing any additional information about the sequence, such as basecalling, alignment, or other metadata. This makes RNAkinet adaptable for future applications since it is independent of external basecaller or alignment software.

### RNAkinet accurately classifies RNA molecules labeled with 5EU

To test the performance of RNAkinet, we first compared the distribution of predicted modification probabilities on the training data. Our results showed that both RNAkinet and RNAkinet-TL could robustly distinguish labeled RNAs and that they were learning the training data (**Sup. Fig. 2A**). We subsequently calculated the receiver operating characteristic curve (ROC) for the testing dataset which consists of reads in chromosome 1, never seen by the NN (**Fig. 2A**). Interestingly, while both models achieved a high area under curve (AUC), RNAkinet performed the best, reaching an AUC of 0.87. In contrast, the model involving transfer learning, RNAkinet-TL, reached an AUC of 0.72. Subsequently we tested the precision and recall (PR) of the two architectures. We again found that RNAkinet outperformed RNAkinet-TL achieving higher precision across the entire dataset with AUC of 0.89 compared to 0.73 (**Fig. 2B**). Since our data can be heavily unbalanced towards the non-labeled class, we calculated the balanced accuracy (BA) as the average of sensitivity and specificity to measure the average accuracy for both the minority and majority classes. Again, RNAkinet outperformed RNAkinet-TL reaching higher average accuracy across all prediction thresholds (**Fig. 2C**). Similar results were also observed for F1 score (**Sup. Fig. 2B**). These data indicate that RNAkinet that involves convolution and recurrent characteristics with only 66,000 parameters, is sufficient to accurately capture 5EU signals in nanopore sequencing reads.

**Figure 2:**
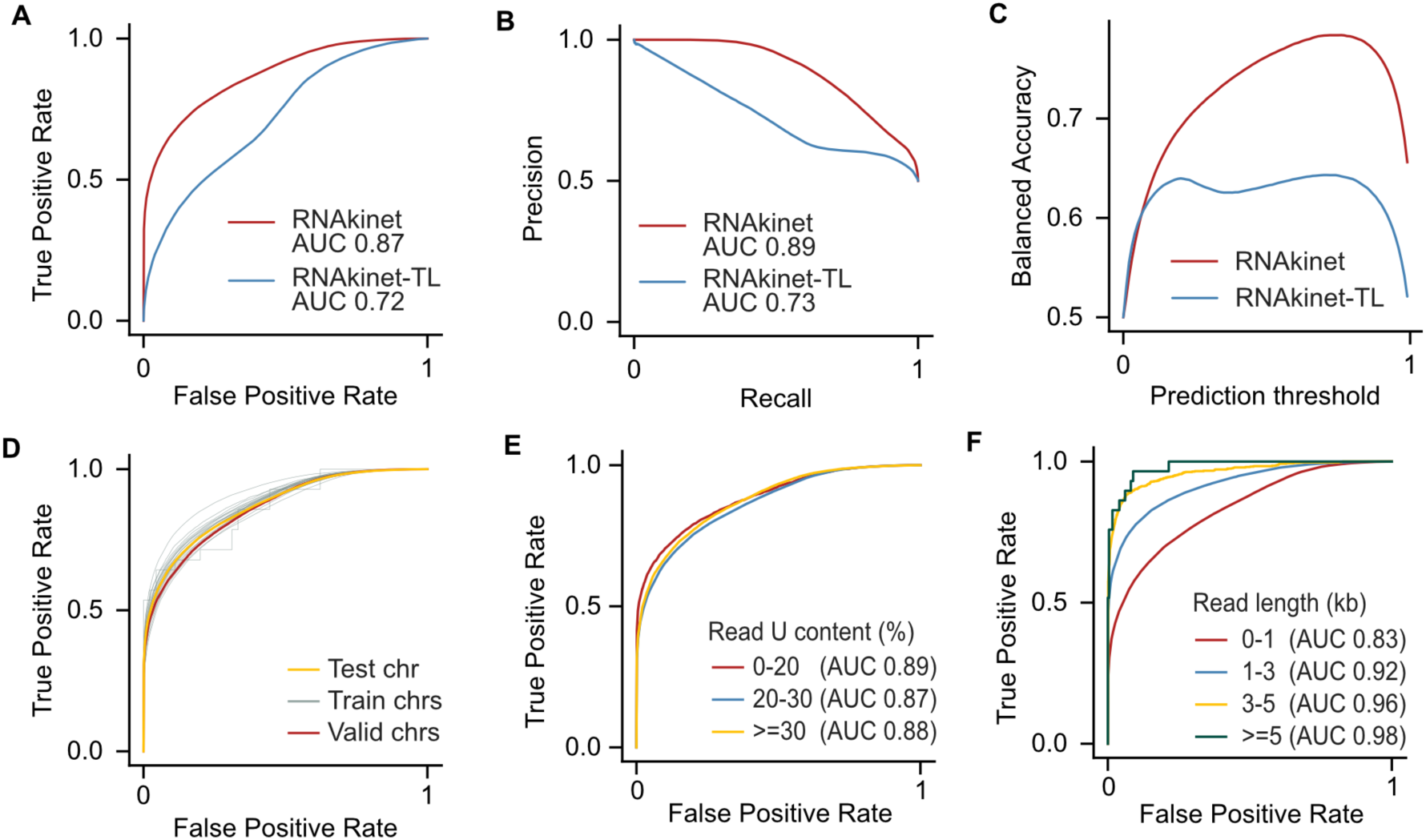
RNAkinet accurately classifies RNA molecules labeled with 5EU. **A-C)** ROC (A), PR (B) and balanced accuracy (C) on test data for RNAkinet and RNAkinet-TL. **D-F)** ROC plot stratified by train, validation, and test chromosomes (D), percent of uridines in transcript (E) and read length (F).

To further explore the training process and validate that RNAkinet does not overfit to reads in the training chromosomes, we calculated the ROC for individual chromosomes and combined **(Fig. 2D, Sup. Fig. 2C-E).** We found that RNAkinet performs similarly across reads from all chromosomes, independently of whether the reads come from training, testing, or validation chromosomes. This indicates that the training process is well controlled, preventing overfitting, and thus increasing the generalization capacity of the trained model. To gain further insight into the model performance on varying read sequences, we stratified our testing reads based on U content and read length. Our data showed that the content of uridines had a negligible impact on performance with the AUC ranging from 0.89 to 0.87 **(Fig. 2E)**. In contrast, the read length appeared to have a larger effect on performance with shorter reads (0-1 kb) reaching AUC of 0.83 compared to higher than 0.92 AUC for longer reads **(Fig. 2F)**. Overall, these results indicate that RNAkinet performance improves with longer reads and is independent of the U content percentage.

### RNAkinet generalizes across cell lines and distinguishes nascent RNA molecules

Our data have indicated that RNAkinet successfully avoids overfitting and we wished to also evaluate whether it can generalize to completely independent datasets. To explore this, we analyzed previously generated nanopore data following labeling of K562 cells with 5EU for 24 h (positive) or control (negative) ^24^. Since this is also a human cell line, we again tested on reads that aligned to chromosome 1 to ensure that RNAkinet had not previously seen the corresponding sequence space and thus performance would not be confounded by read sequence. Our results show that RNAkinet could successfully distinguish labeled RNA molecules along all replicates of K562 cells (**Sup. Fig. 3A**). Quantification of the ROC of the prediction showed that RNAkinet had comparable performance on K562 cells as in HeLa with a drop in performance from 0.87 AUC to 0.80 (**Fig. 3A**). A minor drop in performance was expected given that the data in ^24^ were prepared with an earlier iteration of the ONT RNA sequencing kit (SQK-RNA001) the intricacies of which have not been modeled during training. Similarly, we found that RNAkinet had comparable precision, recall and accuracy on both datasets (**Fig. 3B, Sup. Fig. 3B**). Again, we found that stratification of reads by chromosome, U content or read length resulted in minimal performance differences except for short RNAs (<1kb) that again showed similarly reduced performance as in the original HeLa dataset (**Fig. 3C-E**).

**Figure 3:**
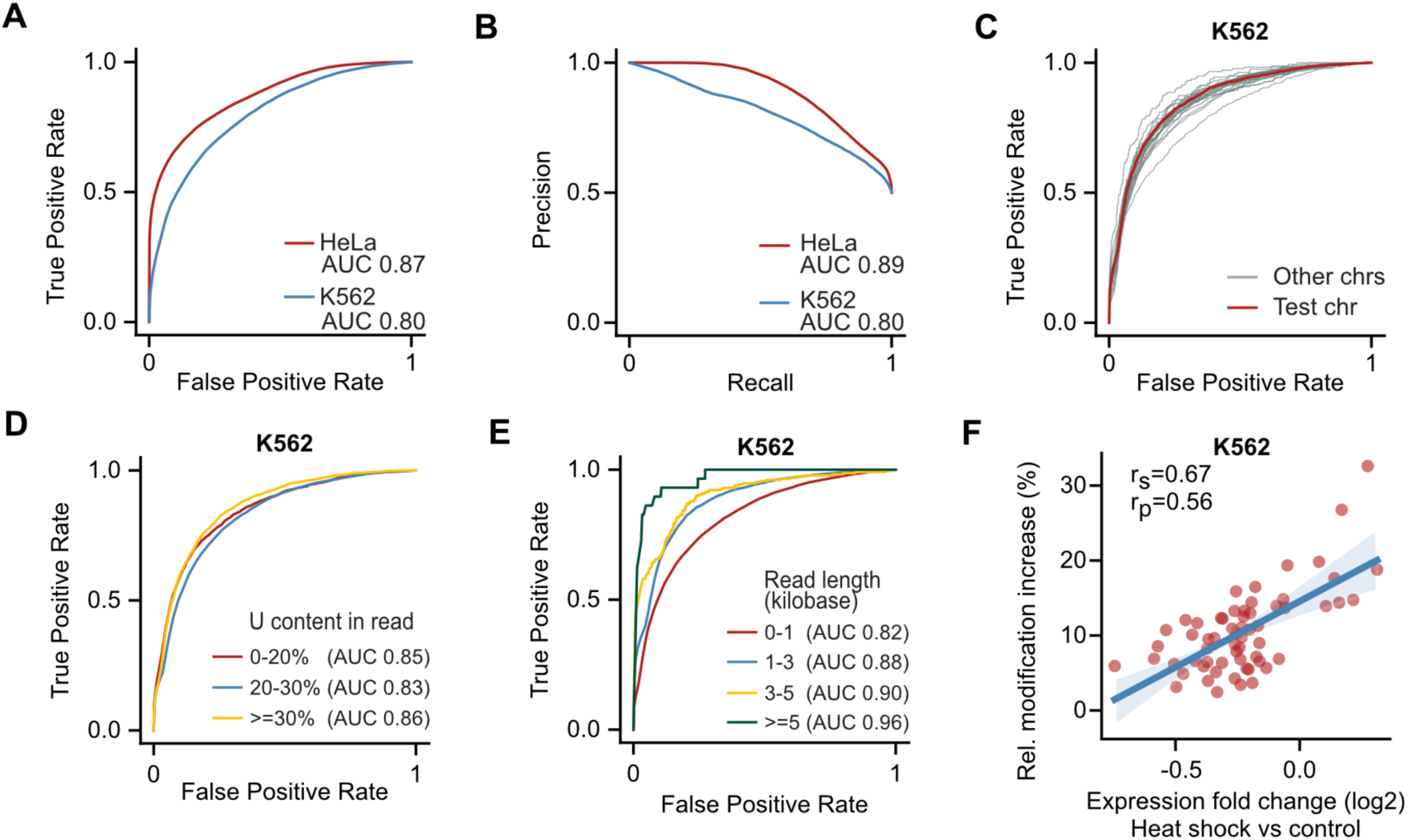
RNAkinet generalizes across cell lines and distinguishes nascent RNA molecules. **A-B)** ROC (A) and PR (B) plot for reads in chromosome 1 for HeLa and K562 cells. Data for HeLa are as in Fig. 2 and only shown here for comparison. **C-E)** ROC plot on K562 cells data stratified by chromosome (C), percent of uridines in transcript (D) and read length (E). **F)** Scatter plot and correlation coefficients of gene expression change and predicted 5EU modification rate for K562 cells subjected to heat shock.

Our results have shown that RNAkinet could successfully identify 5EU-labeled RNA molecules from different cell types in experiments that involve labeling for 24 h. However, such long labeling periods are rarely used in canonical experimental settings. We therefore explored if RNAkinet can be used to predict 5EU-labeled RNAs from shorter labeling periods. We reanalyzed data from ^24^ in which cells were subjected to heat shock at 42°C for 60 min at the presence of 5EU and performed differential expression analysis using DESeq2 ^34^ (**Sup. Fig. 3C**). We reasoned that stress response genes, upregulated upon heat shock, should incorporate more 5EU and thus be recognized as labeled in higher proportions than downregulated genes. We calculated a metric for relative modification increase for genes with at least 100 reads to have substantial read support. As hypothesized, we found that the relative modification increase significantly correlated with expression change, indicating that newly synthesized mRNAs of stress response genes do indeed incorporate 5EU at a higher rate (**Fig. 3F**). Collectively, our results show that RNAkinet can generalize and can accurately identify 5EU-labeled RNA molecules even from short labeling periods and can be used to capture the dynamics of RNA metabolism across conditions.

### RNAkinet predicts RNA isoform kinetics across species

Next, we wished to interrogate the performance of RNAkinet in a different species and to explore whether it can be used to quantify RNA kinetics such as RNA half-life. To test this, we cultured NIH/3T3 cells, a mouse fibroblast cell line in the presence of 5EU for 2 h in biological duplicates. Following labeling, total RNA was harvested and dRNA-Seq was performed. RNAkinet was used to predict 5EU labeling for individual reads and quantified labeled and unlabeled read counts for each gene transcript. Subsequently, we quantified transcript half-lives based on the ratio of modified over unmodified counts (**Fig. 4A**) as previously shown ^35^. Comparison of the two replicates showed high reproducibility with correlation reaching 0.78 for Pearson’s and 0.83 for Spearman’s coefficients (**Fig. 4B**). We re-analyzed half-life measurements produced in ^26^ for the same cell-line using a combination of metabolic labeling with 5EU and isolation azide-bearing biotin tags. Our results show that transcript half-lives quantified by RNAkinet correlated significantly with measured half-lives for both replicates (**Fig. 4C, D**). Moreover, we explored whether read support for each transcript affected transcript half-life predictions. Indeed, we found that higher read support resulted in higher correlation with measured half-lives indicating that higher sequencing depth could further improve performance (**Fig. 3E**). Collectively, our results show that RNAkinet generalizes to RNAs from other species and can quantify RNA kinetics in diverse experimental settings.

**Figure 4:**
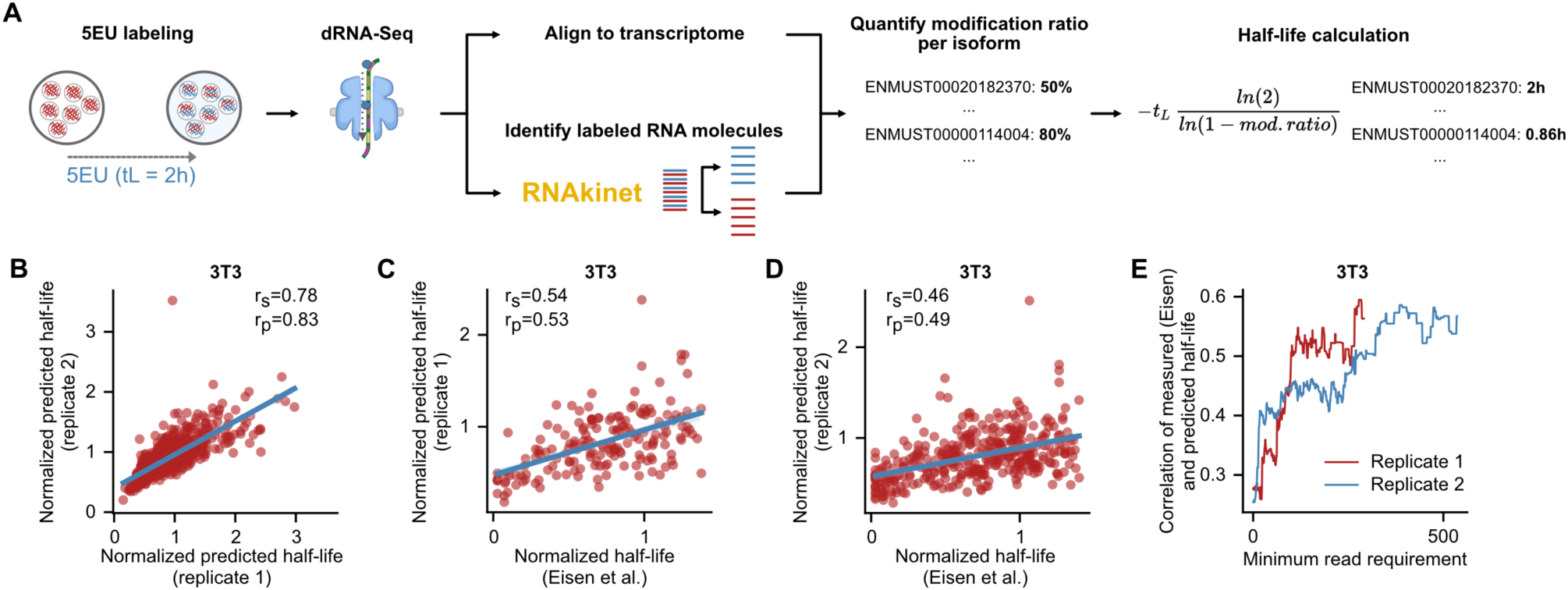
RNAkinet predicts RNA isoform kinetics across species. **A)** Schematic of RNAkinet pipeline to quantify transcript half-lives. **B)** Scatter plot and correlation coefficients of predicted half-lives for 2 independent biological replicates of 3T3 cells. **C-D)** Scatter plot and correlation coefficients of predicted of half-lives quantified in mouse 3T3 cells by ^26^ and RNAkinet for two biological replicates. **E)** Pearson’s correlation coefficient between ^26^ and RNAkinet quantified half-lives for increasing levels of required read coverage per transcript.

## Discussion

The abundance of RNA within cells is determined by a finely controlled balance of synthesis, processing, and degradation, which ensures homeostatic maintenance and responsiveness to environmental signals. Therefore, measuring RNA dynamics is crucial for deciphering cellular regulatory mechanisms in both normal and disease states. In this study, we metabolically labeled newly synthesized RNA with the nucleoside analog 5EU and developed RNAkinet, a tool to detect nascent, labeled RNA molecules directly from raw nanopore signals. To our knowledge, RNAkinet is the only tool that identifies modified nucleotides directly from the raw signal, without dependencies on other software, showing the feasibility of this approach for modification detection. By being able to call the labeling status of individual RNA molecules, RNAkinet offers the unique ability to interrogate a multidimensional ensemble of cis-acting RNA regulatory elements comprising of isoform usage, 5’ end decay, alternative poly(A) site usage, poly(A) tail length and post-transcriptional RNA modifications simultaneously ^12^.

A prevalent limitation of machine learning tools is their potential inability to generalize and extrapolate findings to different systems. To address this, we adopted a thorough data preparation protocol aimed at achieving broad domain representation. Furthermore, we subjected RNAkinet to rigorous validation across a variety of distinct and completely independent experimental conditions, confirming its capability to generalize across various RNA sequencing kits, cell types, and organisms. This also highlights the robustness of the training process which coupled with lack of external dependencies introduces a path to future developments and updates as nanopore sequencing technology advances. We also placed particularly emphasis on model architecture to avoid overparameterization. By incorporating convolution and recurrent layers, RNAkinet captures long-range interactions inherent in the electrical signal while it avoids overfitting to training data thus dramatically reducing the parameter space and achieving high computational efficiency being able to process ∼600,000 reads/h (**Sup. Fig. 4A**).

Finally, we show that RNAkinet can quantify isoform-level RNA kinetic parameters that significantly correlate with measurements from methods that lack isoform-level resolution. This feature enables the study of RNA kinetics at unprecedented resolution allowing associations with post-transcriptional regulatory cues at single molecule resolution. We anticipate that RNAkinet will enable a novel set of study designs that will involve a holistic interrogation of RNA and gene regulation in cells.

## Data availability

Sequencing data have been deposited in the Sequence Read Archive (SRA) under Bioproject accession: PRJNA1030003. The code for the analysis included in the manuscript has been deposited on Zenodo DOI: 10.5281/zenodo.10070389. The code for RNAkinet has been deposited on GitHub (https://github.com/maragkakislab/rnakinet).

## Acknowledgements

We would like to thank Dr. Myong-Hee Sung for her gift of the 3T3 cells and Dr. Michael E. Ward for providing iPSC-derived i3Neurons. This work utilized the computational resources of the NIH HPC Biowulf cluster (http://hpc.nih.gov). Computational resources were also provided by the e-INFRA CZ project (ID:90254), supported by the Ministry of Education, Youth and Sports of the Czech Republic. This research was supported by grand HORIZON-WIDERA-2022 grant BioGeMT (ID: 101086768) to P.A and the Intramural Research Program of the National Institute on Aging, National Institutes of Health grant ZIA AG000696 to MM.

## Author contributions

VM and JM developed RNAkinet and performed computational analysis. CB performed wet-lab experiments with assistance from MJP and SM. VM and MM interpreted the data assisted by JM and PA. VM and MM wrote the manuscript with feedback from all authors.

## Materials and Methods

### Cell culture and metabolic labeling

NIH3T3 cells (ATCC CRL-1658) were cultured at 37°C, 5% CO2, 90% humidity in Dulbecco’s Modified Eagle Medium (Thermo Fisher Scientific) supplemented with 10% bovine calf serum (GeminiBio), 1% MEM-nonessential amino acids (Invitrogen), 2 mM L-Glutamine. Cells were passed weekly by gentle dissociation with trypsin-EDTA 0.25%. HeLa cells (ATCC CCL-2) and HEK-293T (ATCC CRL-3216™) were cultured at 37°C, 5% CO2, 90% humidity in Dulbecco’s Modified Eagle Medium (Thermo Fisher Scientific) supplemented with 10% fetal bovine serum (GeminiBio), 1% MEM-nonessential amino acids (Invitrogen), 2 mM L-Glutamine. iPSC-derived i3Neurons were cultured and differentiated as previously described ^36^. For metabolic labeling, cells were cultured in media containing 400 or 500 μM 5-Ethynyl-uridine (ThermoFisher) for 2 to 24 hours as indicated. Total RNA was extracted using TRIzol reagent (Invitrogen) according to manufacturer’s instructions followed by DNase I treatment (MilliporeSigma). RNA concentration and integrity were determined using a Nanodrop ND-1000 (ThermoFisher) and Qubit^TM^ RNA IQ assay (ThermoFisher), respectively. Library preparation for direct RNA sequencing was performed as previously described ^37^ with modifications. Briefly, poly(A) RNAs were purified from 50 μg of total RNA using Oligo d(T)_25_ Magnetic Beads (New England Biolabs). 500 ng of poly(A) mRNA was used for library preparation using SQK-RNA002 sequencing kit (Oxford Nanopore Technologies). The final library was quantified using Qubit dsDNA High Sensitivity assay kit (ThermoFisher) and sequenced on a MinION device using FLO-MIN106 flow cells (Oxford Nanopore Technologies).

### Data preparation for training, validation, and testing

In order to prevent data leakage and overoptimistic results that could emerge from using k-fold cross-validation with random shuffling, the reads were split into training, validation, and testing sets based on a chromosome they originated from. For evaluating precision, recall, and F1 metrics, which are sensitive to imbalanced data, the splits were up-sampled to have a balanced ratio of positives and negatives. The reads were basecalled with Guppy 6.4.8, aligned to the Ensembl human genome (GRCh38) and transcriptome with Minimap2 ^38^, and then separated into splits based on their mapped chromosome. All reads that did not map to any chromosome and secondary reads were discarded. Chromosome 1 was used for testing, chromosome 20 was used for validation (utilized for early stopping), and the rest were used for training. The first 5000 raw signal values from each read were cropped to avoid sequencing artifacts and any reads shorter than 5000 raw signal values or longer than 400000 raw signal values were discarded. The filtered reads were then normalized by median absolute deviation before being passed through the neural network.

### Neural network design and training

To create a detection tool for 5EU modification, a convolutional neural network classifier was designed. It was trained with data from the training split described above and was optimized to perform a binary classification of raw nanopore signals into either modified or unmodified categories. The network was made up of multiple convolutional blocks extracting local patterns from the input signal, followed by a bidirectional recurrent layer with GRU units that aggregated the extracted information across the whole length of the sequence in both directions. These features were then pooled with max pooling, average pooling, and concatenated with hidden states from the last hidden states of the recurrent layers. To allow the network to accept signals of varying lengths without padding as an input, this pooling layer at the end of the network was utilized to aggregate information across the length dimension into a fixed-size vector. This vector was then fed into a small dense feed-forward network to predict the final 5EU modification score.

For training the network on sequences of variable length, a batch size of size 1 was used. Additionally, to simulate minibatch learning, gradients were accumulated over 64 sequences. The network was trained with a learning rate of 0.001 and weight decay of 0.01 using the AdamW optimizer. The model was trained for 1000 epochs with early stopping on the validation set AUROC metric with threshold 0 and patience of 50 evaluation steps. The first 1000 learning steps were used as warm up steps where the learning rate was scheduled to linearly increase from 0 to the final learning rate of 0.001. To accelerate the network training, a single A100 Nvidia GPU was utilized. The implementation and training of the network were done using PyTorch ^39^ and PyTorch Lightning frameworks. Snakemake ^40^ workflows were developed for the entire process of data splitting, model creation, and evaluation to be reproducible, scalable to large computational clusters, and adaptable to be used on newly sequenced datasets.

### Calculation of relative modification change

To accurately capture the change in gene expression between control and condition experiments using 5EU detection, a metric called relative modification increase was introduced. This metric used the percentage of modified reads of a given transcript from both the condition and control experiments and calculated a relative increase in the condition experiment compared to the control. A positive relative modification increase indicated a higher prevalence of modified reads under the experimental condition, while a negative value indicated a decrease.

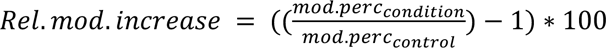, where mod.perc_condition_ and mod.perc_control_ denote the percentage of reads of a given transcript predicted to be modified in the condition and control experiments. These are then divided and 1 is subtracted to represent increase, and result is multiplied by 100 to normalize it to a percentage.

