## Supplementary Tables and Figures for "Deep learning and direct sequencing of labeled RNA captures transcriptome dynamics"

| Dataset | Use case in this work | Generated in | Cell line | Organism |
|---|---|---|---|---|
| In-house HeLa | Positive and negative reads for training, validation, and testing | This work | HeLa | Homo Sapiens |
| In-house Neurons | Negative reads for training | This work | iPSC-derived neurons | Homo Sapiens |
| In-house HEK293T | Negative reads for training | This work | HEK293T | Homo Sapiens |
| In-house 3T3 5EU labeled reads | Testing if model captures RNA decay rates in 3T3 cells | This work | 3T3 | Mus Musculus |
| In-house HeLa 2h 5EU | Testing if model captures rna decay rates in HeLa cells | This work | HeLa | Homo Sapiens |
| Maier K562 classification | Positive (24 hr labeled) reads for testing classification | Maier et al. [9] | K562 | Homo Sapiens |
| Maier K562 classification | Negative (non labeled) reads for testing classification | Maier et al. [9] | K562 | Homo Sapiens |
| Maier K562 heat shock | Test if RNAkinet captures differential expression during stress | Maier et al. [9] | K562 | Homo Sapiens |

**Supplementary Table 1**: Description of dRNA-Seq datasets used in this work.

| **Experiment name** | **Number of reads** |
|---|---|
| hsa_dRNA_HeLa_labeled_1 | 982840 |
| hsa_dRNA_HeLa_nonlabeled_1 | 1878222 |
| mmu_dRNA_3T3_labeled_1 | 620976 |
| mmu_dRNA_3T3_labeled_2 | 1306329 |
| hsa_dRNA_Hek293T_nonlabeled_1 | 1257185 |
| hsa_dRNA_Neuron_nonlabeled_ctrl_1 | 517551 |
| hsa_dRNA_Neuron_nonlabeled_TDP43KD_1 | 487902 |
| hsa_dRNA_HeLa_5EU_2hr_1 | 1911933 |
| hsa_dRNA_HeLa_5EU_2hr_2 | 1174797 |
| hsa_dRNA_HeLa_5EU_2hr_3 | 1364193 |
| 20180514_1054_K562_5EU_1440_labeled_run | 45461 |
| 20180514_1541_K562_5EU_1440_labeled_II_run | 198289 |
| 20180516_1108_K562_5EU_1440_labeled_III_run | 24821 |
| 20180327_1102_K562_5EU_0_unlabeled_run | 144305 |
| 20180403_1102_K562_5EU_0_unlabeled_II_run | 161941 |
| 20180403_1208_K562_5EU_0_unlabeled_III_run | 109778 |

**Supplementary Table 2:** Raw read counts per dataset

**Supplementary Figure 1**

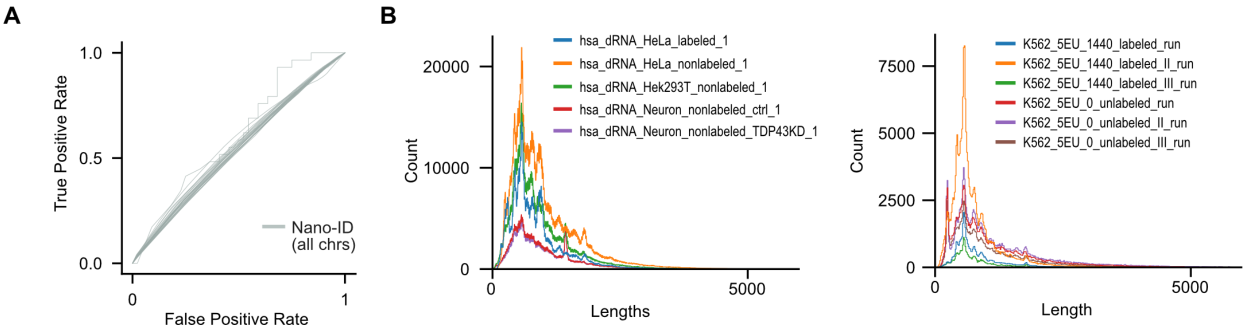


**Supplementary Figure 1: A)** ROC plot of the Nano-ID published in [9] on HeLa cells labeled with 5EU for 24 h. Reads are stratified by chromosome. **B**) Read length distribution for libraries used in training and testing.

**Supplementary Figure 2**

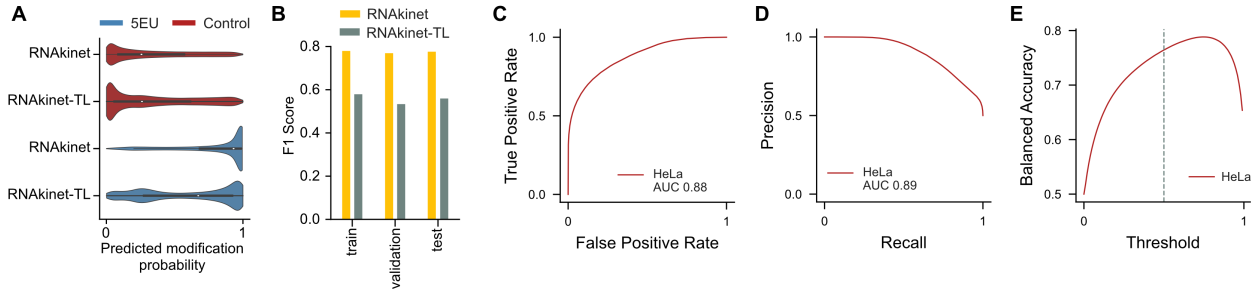


**Supplementary Figure 2: A)** Predicted probability distribution for positive and negative samples used in training for RNAkinet and RNAkinet-TL. **B)** Bar plot of F1 score on train, test, and validation data for RNAkinet and RNAkinet-TL. **C-E)** ROC (C), PR (D) and BA (E) plot for training data for RNAkinet.

### Supplementary Figure 3

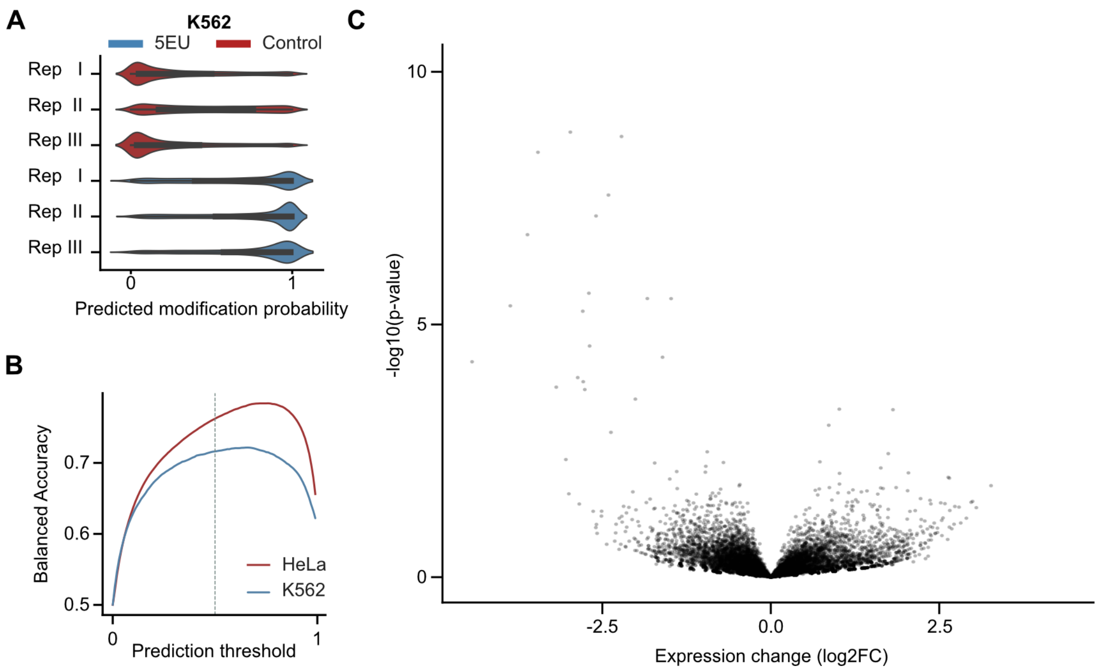


**Supplementary Figure 3: A)** Predicted probability distribution on reads from chromosome 1 for K562 cells **B)** BA on reads from chromosome 1 of HeLa and K562 cells. Data for HeLa cells are the same as Fig. 2 and are only included here for comparison. The threshold used for inference is marked with a gray line. **C)** Volcano plot of isoform differential expression for heat shock against control cells.

### Supplementary Figure 4

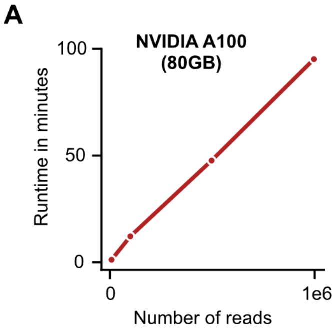


**Supplementary Figure 4: A)** Scatter plot of RNAkinet runtime for inference and number of reads processed.
